# Molecular mechanism decipher of Alzheimer’s disease based on single-cell RNA-seq information

**DOI:** 10.1101/2025.11.22.689872

**Authors:** Xue Jiang, Dongqing Wei, Jing Ye, Eric Gilson

## Abstract

**Background:** Alzheimer’s disease (AD) is a most common age-related neurodegenerative disease with no effective treatments, is one of the leading contributions to global disease burden. AD is represented by complicated biological mechanisms and complexity of brain tissues, often manifested histologically by the parenchymal deposition of amyloid-beta (Aβ) plaques, the formation of neurofibrillary tangles and neuroinflammation. The etiology of AD is likely due to complex interactions among different brain cell types leading to interconnected cellular pathologies. Single-cell RNA sequencing (RNA-seq) technologies enable unbiased characterization of cell types and states, transitions from normal to disease and responses to therapies, providing an effective tool to systematically resolve cellular heterogeneity in AD.

**Methods:** In this study, to investigate the molecular mechanisms of AD, we collected single cell RNA-seq data of disease and control samples and designed an precise computational analysis pipeline for the dataset. We further proposed a consensus non-negative matrix factorization model to extract disease cellular gene expression programs.

**Results:** According to the computational pipeline, we get the most disease-related sub-clusters, cellular process gene expression programs, and disease-related gene programs. We further investigated the enrichment pathways of disease gene programs.

**Discussion:** This research might provide a cell type specific gene program targets therapeutic approach to combat the disease.

**Highlights:**

- We designed a consensus non-negative matrix factorization method, which could extract cellular process gene expression programs, and disease-related gene programs.
- We further investigated the enrichment pathways of the most disease-related sub-clusters.
- We provide a cellular foundation for a new perspective on AD molecular mechanism that informs personalized therapeutic development, targeting the disease-related gene program.

## 1. Background

Alzheimer’s disease is a most common age-related neurodegenerative disease characterized by progressive memory decline and cognitive dysfunction, often manifested histologically by the parenchymal deposition of amyloid-beta (Aβ) plaques, the formation of neurofibrillary tangles and neuroinflammation[1–3]. According to the National Bureau of Health Statistics, AD is the first cause of death among the non-communicable diseases. As there are currently no effective treatments, the Alzheimer’s is one of the leading contributions to global disease burden. Therefore, further efforts should be directed to further understand the molecular mechanism of the disease, prevent the occurrence of the disease or even alleviate the disease through intervention treatment.

Early-onset AD, also known as familial Alzheimer’s disease (fAD), the age of onset is less than 60 years old, accounting for < 5% of the total cases, and the heritability is more than 90%. Late-onset, sporadic Alzheimer’s disease(sAD), the age of onset is more than 60 years old, represents about 95% of all Alzheimer’s disease cases[4], with a complex polygenic background and non-genetic factors, the heritability is 58%-79%. comprehensive and systemic investigations with multi-layered molecular and biological data from different brain regions[5,6], indicated that Alzheimer’s disease is represented by complicated biological mechanisms and complexity of brain tissues. Polygenic risk score studies have found that Alzheimer’s disease involves thousands of genetic risk sites[7]. Meanwhile, deep learning methods, which capture nonlinearity within high-dimensional genomic data, further improve the disease risk prediction [8]. sAD etiology is likely due to complex interactions among different brain cell types leading to interconnected cellular pathologies [9]. It is possible that the cell-independent responses to topologically incorrect protein may be responses to extracellular amyloid deposition, as tau is only intraneuronal and there is evidence of plaque and oligomer interactions with multiple brain cell types. Critically important in the regulation of these processes is the balance between production and clearance of Abeta peptides from the brain. Abeta peptides, the main constituent of senile plaques, are produced mostly by neurons in an activity-dependent manner, and various astrocyte- and microglial-dependent mechanisms are thought to promote breakdown or clearance of Abeta from the brain [10].

Besides, immune cells and pathways are frequently implicated in neurodegenerative conditions. Various innate immune functions have been interchange ably attributed to either microglia or infiltrating blood-derived monocytes. In AD, the local neuroinflammation that is associated with cytotoxicity and disease escalation has often been attributed mainly to microglia[11–14]. At the same time, several studies have also attributed a positive role to infiltrating monocytes in the clearance of toxins from the brain[15–19]. The central nervous system, as an immune privileged site, has evolved unique mechanisms to allow it to benefit from its resident myeloid cells, microglia, as well as from communication with the systemic immune system[20–27].

Due to cellular heterogeneity, defining the roles of immune cell subsets in AD onset and progression has been challenging. Nevertheless, single-cell RNA sequencing (scRNA-seq) technology provides an alternative method to study the cellular heterogeneity of the brain[28–30], by profiling tens of thousands of individual cells[31,32]. Single cell genomic technologies enable unbiased characterization of cell types and states, transitions from normal to disease and response to therapies, supporting comprehensive genome wide sampling by scRNA-seq as an effective tool to systematically resolve immune heterogeneity in AD. Single cell analysis can further identify potential markers, pathways, and regulatory factors, promoting testable hypotheses to elucidate molecular mechanisms of immune regulation in AD[33–35].

In this study, we collected single cell RNA-seq data of disease and control samples of Alzheimer’s disease, and designed a computational analysis pipeline to investigate the cellular mechanism of the disease. We proposed a consensus non-negative matrix factorization model to decompose the gene expression matrix into the product of two lower rank matrices, one is called usage matrix, which encoding the relative contribution of each gene to each program, and the other one is called gene program matrix,which specifying the proportions the programs are combined for each cell. According to the computational framework, we get differentially expressed genes, disease-related sub-cell types, and disease cellular process gene expression programs. We further investigated the enrichment pathways. The computational pipeline help to decipher the cellular mechanisms, providing additional therapeutic avenues aimed at enhancing endogenous cell type-independent responses.

## 2. Methods

### 2.1 Data pre-processing

We denote gene-count matrix as *X*_*ij*_ (*i* = 1,2,…, *N* cells, *j* = 1,2, …, *M* genes). We first select the *H* most over-dispersed genes, which can better distinguish biologically meaningful variations of different samples. The genes was then scaled to unit variance.

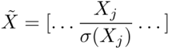 for genes *j* {*H over* − *dispersed genes*} where *X*_*j*_ denotes the *j*-th column of *X, σ* (*X*_*j*_) denotes the sample standard deviation of *X*_*j*_.

The above variance-scaling ensures that genes on different expression scales contribute comparable amounts of information to the gene program inference. Note that we do not perform any cell count normalization (i.e., normalization of the rows of *X*). This is because cells with more counts can contribute more information to the model.

### 2.2 Consensus non-negative matrix factorization (cNMF)

Genes often work together through transcriptional co-regulation, as a gene program in response to the appropriate internal or external signal. We hypothesized that gene programs could be infered directly from single cell expression profiles using matrix factorization. In this context, matrix factorization would model the gene expression matrix as the product of two lower rank matrices, one encoding the relative contribution of each gene to each program, and a second specifying the proportions in which the programs are combined for each cell. We further implemented a meta-analysis approach, which demonstrably increased robustness and accuracy. Overall, the meta-analysis of NMF, which we call consensus NMF (cNMF).

We extract the usage matrix and program matrix by the decomposition of non-negative matrix factorization(NMF). *R* replicates of NMF are run on the same normalized dataset with the same number of components *K* but with different randomly selected seeds, resulting in *R* instances of usage matrices *U*^(*r*)^(*N cells* × *K programs*) and program matrices *G*^(*r*)^ (*K programs* × *g cells*). The Schematics of the computational pipeline is shown in Figure 1.

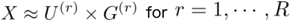

**Figure 1.**
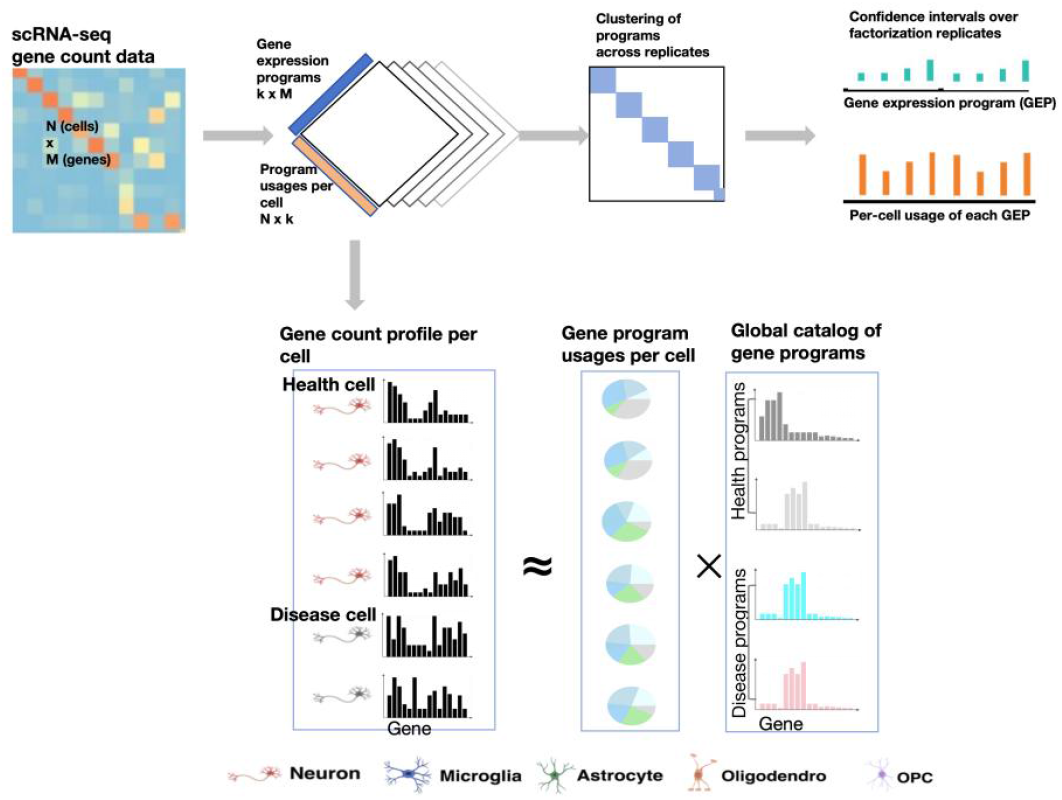
Schematics of of the consensus matrix factorization pipeline and the matrix factorization for single-cell RNA-seq analysis. The gene expression profiles of individual cells are weighted mixtures of a set of global gene expression programs with distinct weights reflecting the usage of each program.

For each replicate *r*, the rows of *G*^(*r*)^ are normalized to have *l*2 norm of 1:

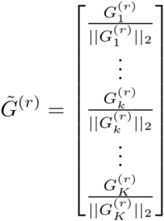

where 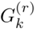 is *k* − *th* the row of the programs matrix for the NMF replicate *G*^(*r*)^, and ‖ ‖_2_ denotes the *l*2 norm. The component matrices from each replicate are then concatenated vertically into a single *RK* × *H* dimensional matrix, *G*,where each row is a component from one replicate:

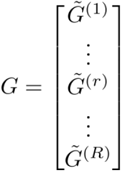

Components with great mean Euclidean distance from their *L* nearest neighbors are then filtered out as below:

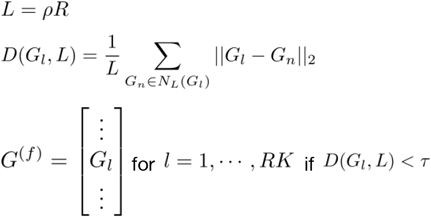

where *G*_*l*_ is the *l* − *th* row of *G, N*_*l*_ (*G*_*l*_) is the set of nearest neighbors of and is the matrix of rows that passed the *L* nearest neighbors distance threshold filter.

Two user-specified parameters, *ρ* and *τ*,determine which replicate components are filtered out and which are kept. *ρ* denotes the fraction of NMF replicates to be used as nearest neighbors. Intuitively, *ρ* can be thought of as the fraction of replicates that must yield a component approximately matching a program in order that program to be kept. *τ* is a distance threshold that determines how close a component must be to its neighbors in Euclidean space to be considered ‘approximately matching’.

We set *ρ* = 0.3 for all datasets analyzed in this manuscript which reflects a tolerance to identify components that occur approximately in 30% or more replicates.

Next, the rows of *G*^(*f)*^ are clustered using K-Means with the Euclidean distance metric and the same number of clusters (K) as the number of components for the NMF runs. This defines sets *A*_*k*_ {*rows assigned to cluster k*} containing the indices of *G*^(*f)*^ the rows of that are assigned to the *k* − *th* cluster. Each cluster of replicate components is then collapsed down to a single consensus vector by taking the median value for each gene across components in a cluster:

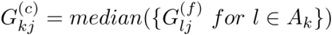

This defines a *K*×*H* consensus programs matrix *G*^(*c)*^ where the (*C*) superscript denotes consensus. The merged gene program components are then *l*1 normalized:

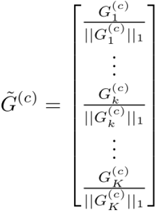

where 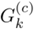 is the *k* − *th* row of *G*^(*c)*^ and ‖ ‖_1_ denotes the *l*1 norm. A consensus usage matrix is then fit by running one last iteration of NMF with the program matrix fixed to 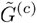.

This amounts to fitting non-negative least squares regression of each cell’s normalized expression profile 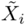 against 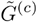 by solving the following optimization

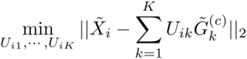

where 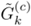 is the *H*-dimensional normalized consensus program vector for the *k* − *th* gene program, *i* indexes over cells, and we are maximizing with respect to gene program usage values 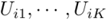 which are constrained to be non-negative. We concatenate all these coefficients into a consensus usage matrix *U*(*c*):

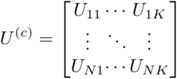

and normalize it so that the usage values for each cell sum to 1:

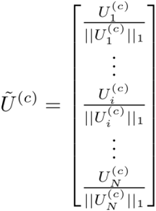

where 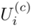 is the *i* − *th* row of the consensus usage matrix. With this normalized, consensus usage matrix fixed, final program estimates can be computed for all genes, including genes that were not initially included among the over-dispersed set. This is done by running a last iteration of NMF with the usage matrix fixed as 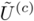 and the input data reflecting the desired genes.

### 2.3 Cellular process gene expression programs

For the scRNA-seq data of disease-related sub-clusters, we extracted gene expression programs that are involved in the disease process with the cNMF. Let *X*_*ij*_ (*i* = 1,2,…, *N* cells, *j* = 1,2, …, *M* genes)be the observed gene-count matrix of disease-related sub-clusters. We assume the cNMF for *X*_*ij*_ as follows:

Following the multiplicative update rules of cNMF, the convergence has been achieved after 100 iterations or when the reconstruction error is below a user-specified error threshold (here the threshold is taken to be 1 × 10^−4^).

For the dataset, we specified the number of gene expression programs (latent factor) *K* to balance the reconstruction stability and error. For each latent factor, we define a cellular process gene program by identifying genes with high correlation (across cells) between expression in a cell and the contribution of each factor to each cell. We annotate each cellular process gene expression program by the pathways most enriched in the genes.

### 2.4 Marker genes identification

The main cell types of the human brain include neurons, inhibitory neurons, excitatory neurons, oligodendrocytes, oligodendrocyte progenitor cells, microglia, astrocytes, endothelial cells, and pericytes. Marker genes for the major cell types of the human brain were obtained from previously published studies [36–40]. neurons marked by SYT1, SNAP25, GRIN1, GRIK2, GRIA1, GRIN2B, and RBFOX1. Inhibitory neurons marked by GAD1 and GAD2. Excitatory neurons marked by SLC17A7, CAMK2A, and NRGN. Oligodendrocytes marked by MBP, MOBP, and PLP1. Oligodendrocyte progenitor cells marked by PDGFRA, VCAN, CSPG4, PCDH15, and MEGF11. Microglia marked by CD74, CSF1R, HLA-DRA, CX3CR1, C1QB, CSF1R, and KCNQ3. Astrocytes marked by AQP4, GFAP, SLC1A2, and ADGRV1. Endothelial cell marked by FLT1 and CLDN5. Pericytes marked by AMBP.

Brain cell-type marker gene sets curated from independent human single-cell RNA datasets for cell type proportion estimation in bulk RNA datasets. For each pre-cluster, we assigned a cell-type label using statistical enrichment for sets of marker genes, and manual evaluation of gene expression for small sets of known marker genes. Enrichment was statistically assessed using the hypergeometric distribution (Fisher’s exact test) and FDR correction over all gene sets and pre-clusters. Broad cell-type clusters were defined by grouping together all pre-clusters corresponding to the same cell type. Sub-clustering analysis was performed independently over each broad cell-type cluster.

### 2.5 Gene module and pathway enrichment analysis

For each gene expression program, according to gene-level significance score, we considered the top 200 ranked genes for further enrichment analysis. Gene module and pathway enrichment analysis was done using Metascape (http://metascape.org/) online tool with default parameters [41].

## 3. Results

### 3.1 Sample preparation and data description

We collected datasets from the research of Mathys H, et al [42] and Grubman A, et al [43]. The detailed information of participant demographics and characteristics are shown in Table 1. The Grubman A dataset applied single-nucleus RNA sequencing to entorhinal cortex samples of 6 control and 6 Alzheimer’s disease brains. The raw expression matrix was composed of 33,694 genes and 14,876 cells. The Mathys H, et al applied single-nucleus RNA sequencing to prefrontal cortex samples of 48 individuals with varying degrees of Alzheimer’s disease pathology (24 no pathology and 24 AD-pathology). The raw expression matrix was composed of 80,660 cells, with a median value of 1,496 total read counts over protein-coding genes. The preprocess steps for the two dataset are shown below respectively.

**Table 1.** Participant demographics and characteristics.

| Sample | Cell number | Sex | Age | PMI | Braak Staging | COGDX | CDX | AD or Not | Artical |
| --- | --- | --- | --- | --- | --- | --- | --- | --- | --- |
| ROS1 | 2913 | Female | 90+ | 4.166667 | 1 | 1 | 1 | NO | [1] |
| ROS2 | 1401 | Female | 80.1807 | 15.33333 | 3 | 1 | 1 | NO | [1] |
| ROS3 | 1329 | Female | 83.72895 | 18.21667 | 3 | 1 | 1 | NO | [1] |
| ROS4 | 1230 | Female | 79.53457 | 4.5 | 2 | 4 | 4 | NO | [1] |
| ROS5 | 1749 | Female | 90+ | 7 | 4 | 1 | 1 | NO | [1] |
| ROS6 | 745 | Female | 79.11567 | 5.333333 | 3 | 6 | 5 | NO | [1] |
| ROS7 | 937 | Female | 87.30185 | 3.25 | 4 | 1 | 1 | NO | [1] |
| ROS8 | 1171 | Female | 76.05749 | 3.583333 | 3 | 5 | 4 | NO | [1] |
| ROS9 | 839 | Female | 87.66598 | 16.71667 | 3 | 2 | 2 | NO | [1] |
| ROS10 | 1809 | Female | 88.92539 | 6.283333 | 1 | 2 | 2 | NO | [1] |
| ROS11 | 1533 | Female | 74.77344 | 3.916667 | 3 | 2 | 2 | NO | [1] |
| ROS12 | 1699 | Female | 90+ | 20.83333 | 3 | 3 | 3 | NO | [1] |
| ROS13 | 484 | Male | 87.23888 | 1.333333 | 3 | 2 | 2 | NO | [1] |
| ROS14 | 1169 | Male | 86.91581 | 8.583333 | 3 | 1 | 1 | NO | [1] |
| ROS15 | 2041 | Male | 89.71389 | 16.75 | 2 | 1 | 1 | NO | [1] |
| ROS16 | 1761 | Male | 80.53388 | 15.16667 | 1 | 1 | 1 | NO | [1] |
| ROS17 | 3364 | Male | 89.02669 | 3.25 | 2 | 2 | 2 | NO | [1] |
| ROS18 | 1556 | Male | 90+ | 2.583333 | 2 | 1 | 1 | NO | [1] |
| ROS19 | 1424 | Male | 90+ | 16.66667 | 3 | 1 | 1 | NO | [1] |
| ROS20 | 308 | Male | 90+ | 22.33333 | 2 | 4 | 6 | NO | [1] |
| ROS21 | 815 | Male | 84.98836 | 6.5 | 3 | 4 | 4 | NO | [1] |
| ROS22 | 1706 | Male | 80.94456 | 7.583333 | 3 | 2 | 2 | NO | [1] |
| ROS23 | 1710 | Male | 90+ | 85.08333 | 2 | 1 | 1 | NO | [1] |
| ROS24 | 1447 | Male | 88.46817 | 2.333333 | 1 | 3 | 2 | NO | [1] |
| ROS25 | 1558 | Female | 86.26968 | 3.25 | 5 | 4 | 4 | YES | [1] |
| ROS26 | 2279 | Female | 83.49624 | 4.25 | 5 | 4 | 4 | YES | [1] |
| ROS27 | 1299 | Female | 87.44969 | 18.16667 | 4 | 4 | 5 | YES | [1] |
| ROS28 | 1855 | Female | 83.72895 | 26.16667 | 5 | 4 | 4 | YES | [1] |
| ROS29 | 764 | Female | 76.88433 | 14 | 5 | 3 | 2 | YES | [1] |
| ROS30 | 1023 | Female | 90+ |  | 3 | 2 | 2 | YES | [1] |
| ROS31 | 976 | Female | 83.39767 | 2.166667 | 5 | 4 | 4 | YES | [1] |
| ROS32 | 2853 | Female | 87.50992 | 4.25 | 5 | 4 | 4 | YES | [1] |
| ROS33 | 1521 | Female | 90+ | 7.75 | 5 | 4 | 5 | YES | [1] |
| ROS34 | 1572 | Female | 87.95346 | 5.083333 | 5 | 4 | 4 | YES | [1] |
| ROS35 | 1877 | Female | 86.13005 | 16.68333 | 6 | 4 | 4 | YES | [1] |
| ROS36 | 2315 | Female | 82.71047 | 16.93333 | 5 | 4 | 4 | YES | [1] |
| ROS37 | 2771 | Male | 80.09856 | 20.48333 | 5 | 1 | 1 | YES | [1] |
| ROS38 | 2024 | Male | 85.82888 | 17.91667 | 3 | 4 | 4 | YES | [1] |
| ROS39 | 1111 | Male | 86.93224 | 13.25 | 5 | 4 | 4 | YES | [1] |
| ROS40 | 350 | Male | 81.67556 | 1.5 | 3 | 4 | 4 | YES | [1] |
| ROS41 | 1490 | Male | 90+ | 4.5 | 5 | 4 | 5 | YES | [1] |
| ROS42 | 1274 | Male | 81.46749 | 4.5 | 4 | 4 | 4 | YES | [1] |
| ROS43 | 1550 | Male | 90+ | 4.25 | 4 | 1 | 1 | YES | [1] |
| ROS44 | 713 | Male | 83.63587 | 4.583333 | 5 | 2 | 2 | YES | [1] |
| ROS45 | 297 | Male | 85.77687 | 3.5 | 6 | 4 | 4 | YES | [1] |
| ROS46 | 1304 | Male | 86.40383 | 13.83333 | 6 | 4 | 4 | YES | [1] |
| ROS47 | 1516 | Male | 90+ | 8.666667 | 5 | 4 | 4 | YES | [1] |
| ROS48 | 1202 | Male | 88.39973 | 17.75 | 4 | 1 | 1 | YES | [1] |
| AD1 | 1554 | Male | 91 | 8 | 6 | -- | -- | YES | [2] |
| AD2 | 1426 | Male | 83.8 | 10 | 6 | -- | -- | YES | [2] |
| AD3 | 1243 | Female | 67.8 | 21 | 6 | -- | -- | YES | [2] |
| AD4 | 1342 | Female | 83 | 34 | 6 | -- | -- | YES | [2] |
| AD5 | 484 | Male | 73 | 9.5 | 6 | -- | -- | YES | [2] |
| AD6 | 439 | Male | 74.6 | 30 | 5 | -- | -- | YES | [2] |
| Ct1 | 750 | Female | 67.3 | 24 | -- | -- | -- | NO | [2] |
| Ct2 | 305 | Female | 82.7 | 28.5 | -- | -- | -- | NO | [2] |
| Ct3 | 1964 | Male | 72.6 | 42.5 | -- | -- | -- | NO | [2] |
| Ct4 | 1006 | Male | 75.6 | 46 | -- | -- | -- | NO | [2] |
| Ct5 | 1191 | Male | 77.5 | 53.5 | -- | -- | -- | NO | [2] |
| Ct6 | 1066 | Male | 82.7 | 27 | -- | -- | -- | NO | [2] |

MMSE: mini-mental state examination
PMI: postmortem interval
ANP: assessment of neuritic plaques
COGDX: final clinical dx-clinical consensus diagnosis
CDS: clinical cognitive diagnosis summary at last visit
Braak staging[1-3]: a system for staging the severity of Alzheimer's disease (AD), based on the premise that AD pathology—specifically neurofibrillary tangles—evolve in stages and locations, beginning in the mesial temporal lobe (stage 1 and 2) and extending to the limbic regions (stage 3 and 4) – at which point dementia manifests itself clinically ending in the neocortex.

The preprocess for Grubman A dataset:

Step1. genes without any counts in any cells were filtered out. Besides, the 100 postmortem interval (PMI)-associated genes detected in the data were removed [44].

Step2. cells outside the 5th and 95th percentile with respect to the number of genes detected and the number of minimum unique molecular identifiers (UMIs) were discarded. In addition, cells with more than 10% of their UMIs assigned to mitochondrial genes were filtered out.

Step3. the matrix was normalized with a scale factor of 1000 as recommended by the Seurat pipeline [45] with the parameters x.low.cutoff = 0.0125, x.high.cutoff = 3, and y.cutoff=0.5. overall, the resulting filtered matrix consisted of 10850 genes and 13214 high-quality nuclei.

The preprocess for Mathys H dataset:

To control the data quality for downstream analysis, following steps were conducted:

Step1. only counts associated with protein-coding genes were considered; mitochondrially encoded genes and genes detected in fewer than 2 cells were excluded.

Step2. cells with fewer than 200 detected genes and cells with an abnormally high ratio of counts mapping to mitochondrial genes (relative to the total number of detected genes) were removed.

Step3. normalization and clustering were done with the SCANPY package. The dataset was projected onto the two-dimensional space using t-distributed stochastic neighbour embedding (t-SNE) on the top 10 principal components.

Step4. pre-clusters of 360 and 791 cells that reflected low-quality cells were excluded (that is, cells that showed mixed cell-type markers; extreme complexity with many more genes expressed than other cells; either too many or too few reads; and, in one case, cells isolated almost exclusively from one individual).

Step5. for each pre-cluster, differentially expressed genes were detected using the variance-adjusted t-test as implemented in the function rank_genes_groups in SCANPY. The top 500 ranking genes were extracted for each cluster, and used to test for overlap with markers as previously reported. Those having a distinctly high number of total counts and mixed expression of markers from different cell types were tagged as potential doublets and not considered for downstream analyses.

After applying these filtering steps, a gene-count matrix with 17,926 genes and 70,634 cells were generated. Finally, the 10,850 genes and 13,214 cells of Grubman dataset, and the 17,926 genes and 70,634 cells were combined into a single dataset, by merging all the cells with same genes. Finally, we get 9857 genes and 83,848 high-quality cells for downstream analysis. The number of filtered cells for the participants are shown in Table 2.

**Table 2.** The number of filtered cells for the participants.

| **Sample** | **AD or Not** | **Cell number** |
| --- | --- | --- |
| AD1-6 | AD | 6673 |
| Ct1-6 | no-pathology | 6541 |
| ROS1 | no-pathology | 2893 |
| ROS2 | no-pathology | 1394 |
| ROS3 | no-pathology | 1321 |
| ROS4 | no-pathology | 1221 |
| ROS5 | no-pathology | 1741 |
| ROS6 | no-pathology | 739 |
| ROS7 | no-pathology | 934 |
| ROS8 | no-pathology | 1166 |
| ROS9 | no-pathology | 836 |
| ROS10 | no-pathology | 1796 |
| ROS11 | no-pathology | 1523 |
| ROS12 | no-pathology | 1689 |
| ROS13 | no-pathology | 476 |
| ROS14 | no-pathology | 1156 |
| ROS15 | no-pathology | 2023 |
| ROS16 | no-pathology | 1745 |
| ROS17 | no-pathology | 3349 |
| ROS18 | no-pathology | 1548 |
| ROS19 | no-pathology | 1412 |
| ROS20 | no-pathology | 307 |
| ROS21 | no-pathology | 809 |
| ROS22 | no-pathology | 1702 |
| ROS23 | no-pathology | 1698 |
| ROS24 | no-pathology | 1416 |
| ROS25 | early-pathology | 1544 |
| ROS26 | late-pathology | 2264 |
| ROS27 | early-pathology | 1286 |
| ROS28 | early-pathology | 1841 |
| ROS29 | early-pathology | 761 |
| ROS30 | early-pathology | 1018 |
| ROS31 | late-pathology | 963 |
| ROS32 | early-pathology | 2834 |
| ROS33 | early-pathology | 1514 |
| ROS34 | late-pathology | 1562 |
| ROS35 | late-pathology | 1865 |
| ROS36 | early-pathology | 2304 |
| ROS37 | early-pathology | 2743 |
| ROS38 | early-pathology | 1998 |
| ROS39 | late-pathology | 1106 |
| ROS40 | early-pathology | 346 |
| ROS41 | late-pathology | 1486 |
| ROS42 | early-pathology | 1271 |
| ROS43 | early-pathology | 1539 |
| ROS44 | early-pathology | 709 |
| ROS45 | late-pathology | 287 |
| ROS46 | late-pathology | 1298 |
| ROS47 | late-pathology | 1500 |
| ROS48 | early-pathology | 1192 |

### 3.2 Cell type annotation and sub-clustering

Normalization and clustering were done with the SCANPY package. In brief, counts for all nuclei were scaled by the total library size multiplied by 10,000 and transformed to log space. A total of 2,186 highly variable genes were identified based on dispersion and mean, the technical influence of the total number of counts was regressed out, and the values were rescaled. These preprocessing steps were performed by sequentially using the functions normalize_total(), highly_variable_genes(), regress_out(), scale() in SCANPY. 509 cells that have less than 200 genes expressed were filtered out.

Principal component analysis (PCA) was performed on the left cells with the variable genes, and t-SNE was run on the top 10 principal components (PCs) using the Multicore-TSNE package (https://github.com/DmitryUlyanov/Multicore-TSNE). The top 50 PCs were used to build a k-nearest-neighbours cell-cell graph with k=30 neighbours;and the Louvain graph-clustering algorithm was applied to identify cell clusters. These analyses were performed using the functions pca, neighbours, and louvain in SCANPY.

Visualization of single-cell transcriptomes in uniform manifold approximation and projection (UMAP) space was able to separate cell into clusters, which we mapped to the eight cell types, including neurons, inhibitory neurons (Inh.), excitatory neurons (Exc.), oligodendrocytes (Oli.), oligodendrocyte progenitor cells (OPC), microglia (Mic.), astrocytes (Ast.), and endothelial cells(End.), based on previously established marker gene sets. The initial pre-clustering analysis resulted in 24 pre-clusters with a median number of 2590 cells, ranging from 174 to 11,326 cells. The number of pre-clusters for each cell type are as follows: all cell types, 24; neurons, 13; inhibitory neurons, 4; excitatory neurons, 9; oligodendrocytes, 6; oligodendrocyte progenitor cells, 1; microglia, 1; astrocytes, 1; endothelial cells, 1; unknown, 1. The results are shown in Figure 2.

**Figure 2.**
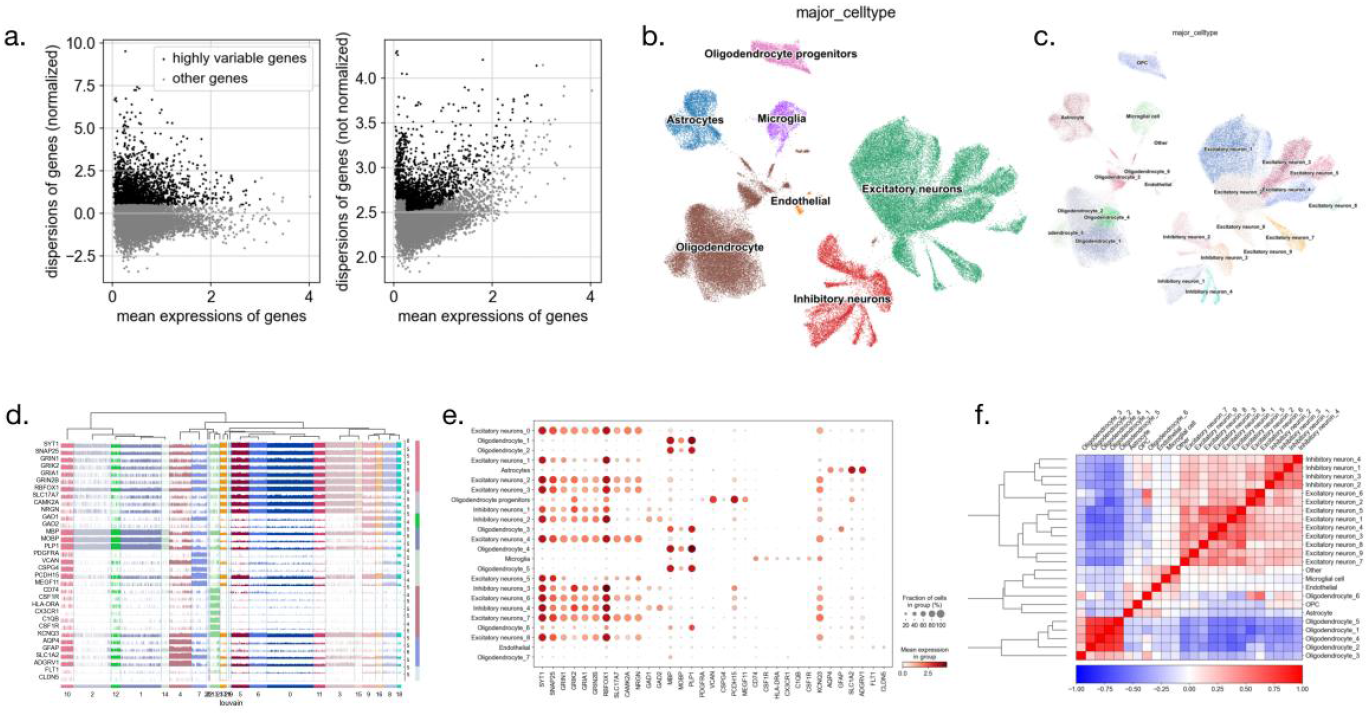
Results for cell type annotation and sub-clustering. (a) Volcano diagram of highly variable genes. (b) cell type clustering and annotation. (c) sub-cell type clustering and annotation. (d) the hierarchical clustering with biomarker gene expression in each cell type. (e) the expression of marker gene in each cell type. (f) heatmap of correlations between different sub-cell types.

For each pre-cluster, we assigned a cell-type label using statistical enrichment for sets of marker genes, and manual evaluation of gene expression for small sets of known marker genes. Sub-clustering analysis was performed independently over each broad cell-type cluster.

Next,we conducted detailed statistics of cell proportions between disease samples and normal control samples in each sub-cluster. The number of cells for each cell sub-cluster is shown in Table 3. The proportion of cells for each cell sub-cluster is shown in Table 4. We found that in entorhinal cortex the amount of astrocytes and oligodendrocyte are significantly decreased, the amount of inhibitory neurons and endothelial are slightly decreased, while the amount of oligodendrocyte are slightly increased. These results indicated that the disease-related sub-cell types include Exc.2, Ast, OPC, Oli.3, Oli.4, Oli.5, Oli.6, and End.

**Table 3.** The number of cells for each cluster(cell sub-population.

|  |  | Entorhinal Cortex |  | Prefrontal Cortex |  |  |
| --- | --- | --- | --- | --- | --- | --- |
|  |  | no-patholog<br>y(6) | AD(6) | no-patholog<br>y(24) | early-pathol<br>ogy(15) | late-patholo<br>gy(9) |
| Cluster_0 | Excitatory neurons_1 | 6 | 4 | 6475 | 3065 | 1776 |
| Cluster_1 | Oligodendrocyte_1 | 227 | 229 | 4498 | 2970 | 2068 |
| Cluster_2 | Oligodendrocyte_2 | 426 | 440 | 4523 | 2522 | 1265 |
| Cluster_3 | Excitatory neurons_2 | 16 | 19 | 2282 | 3601 | 1281 |
| Cluster_4 | Astrocytes | 1719 | 384 | 1553 | 1270 | 543 |
| Cluster_5 | Excitatory neurons_3 | 2 | 1 | 2305 | 1439 | 689 |
| Cluster_6 | Excitatory neurons_4 | 1 | 4 | 2208 | 1431 | 734 |
| Cluster_7 | Oligodendrocyte<br>progenitors | 957 | 179 | 1335 | 867 | 419 |
| Cluster_8 | Inhibitory neurons_1 | 182 | 51 | 1608 | 1110 | 466 |
| Cluster_9 | Inhibitory neurons_2 | 23 | 20 | 1591 | 890 | 656 |
| Cluster_10 | Oligodendrocyte_3 | 16 | 3075 | 1 | 1 | -- |
| Cluster_11 | Excitatory neurons_5 | -- | -- | 1393 | 949 | 546 |
| Cluster_12 | Oligodendrocyte_4 | 2061 | 39 | 85 | 43 | 64 |
| Cluster_13 | Microglia | 316 | 229 | 835 | 465 | 396 |
| Cluster_14 | Oligodendrocyte_5 | 4 | 1889 | 21 | 22 | 13 |
| Cluster_15 | Excitatory neurons_6 | 9 | 3 | 1005 | 434 | 421 |
| Cluster_16 | Inhibitory neurons_3 | 48 | 12 | 1060 | 444 | 306 |
| Cluster_17 | Excitatory neurons_7 | 1 | 3 | 780 | 581 | 277 |
| Cluster_18 | Inhibitory neurons_4 | 36 | 33 | 559 | 320 | 137 |
| Cluster_19 | Excitatory neurons_8 | 1 | 0 | 313 | 230 | 122 |
| Cluster_20 | Oligodendrocyte_6 | 408 | 10 | 1 | -- | -- |
| Cluster_21 | Excitatory neurons_9 | 1 | -- | 211 | 133 | 70 |
| Cluster_22 | Endothelial | 79 | 44 | 148 | 74 | 58 |
| cluster_23 | Other | 2 | 5 | 104 | 39 | 24 |
| total |  | 6541 | 6673 | 34894 | 22900 | 12331 |

**Table 4.** The proportion of cells for the sample in each cluster(cell sub-population.

|  |  | Entorhinal Cortex |  |  |  | Prefrontal Cortex |  |  |  |  |  |
| --- | --- | --- | --- | --- | --- | --- | --- | --- | --- | --- | --- |
|  |  | no-pathology(6) |  | AD(6) |  | no-pathology(24) |  | early-pathology(15) |  | late-pathology(9) |  |
| Cluster_0 | Excitatory neurons_1 | 6 | 0.0009 | 4 | 0.0006 | 6475 | 0.1856 | 3065 | 0.1338 | 1776 | 0.1440 |
| Cluster_1 | Oligodendrocyte_1 | 227 | 0.0347 | 229 | 0.0343 | 4498 | 0.1289 | 2970 | 0.1297 | 2068 | 0.1677 |
| Cluster_2 | Oligodendrocyte_2 | 426 | 0.0651 | 440 | 0.0659 | 4523 | 0.1296 | 2522 | 0.1101 | 1265 | 0.1026 |
| Cluster_3 | Excitatory neurons_2 | 16 | 0.0024 | 19 | 0.0028 | 2282 | 0.0654 | 3601 | 0.1572 | 1281 | 0.1039 |
| Cluster_4 | Astrocytes | 1719 | 0.2628 | 384 | 0.0575 | 1553 | 0.0445 | 1270 | 0.0555 | 543 | 0.0440 |
| Cluster_5 | Excitatory neurons_3 | 2 | 0.0003 | 1 | 0.0001 | 2305 | 0.0661 | 1439 | 0.0628 | 689 | 0.0559 |
| Cluster_6 | Excitatory neurons_4 | 1 | 0.0002 | 4 | 0.0006 | 2208 | 0.0633 | 1431 | 0.0625 | 734 | 0.0595 |
| Cluster_7 | Oligodendrocyte progenitors | 957 | 0.1463 | 179 | 0.0268 | 1335 | 0.0383 | 867 | 0.0379 | 419 | 0.0340 |
| Cluster_8 | Inhibitory neurons_1 | 182 | 0.0278 | 51 | 0.0076 | 1608 | 0.0461 | 1110 | 0.0485 | 466 | 0.0378 |
| Cluster_9 | Inhibitory neurons_2 | 23 | 0.0035 | 20 | 0.0030 | 1591 | 0.0456 | 890 | 0.0389 | 656 | 0.0532 |
| Cluster_10 | Oligodendrocyte_3 | 16 | 0.0024 | 3075 | 0.4608 | 1 | 0.00003 | 1 | 0.00004 | -- | -- |
| Cluster_11 | Excitatory neurons_5 | -- | -- | -- | -- | 1393 | 0.0399 | 949 | 0.0414 | 546 | 0.0443 |
| Cluster_12 | Oligodendrocyte_4 | 2061 | 0.3151 | 39 | 0.0058 | 85 | 0.0024 | 43 | 0.0019 | 64 | 0.0052 |
| Cluster_13 | Microglia | 316 | 0.0483 | 229 | 0.0343 | 835 | 0.0239 | 465 | 0.0203 | 396 | 0.0321 |
| Cluster_14 | Oligodendrocyte_5 | 4 | 0.0006 | 1889 | 0.2831 | 21 | 0.0006 | 22 | 0.0010 | 13 | 0.0011 |
| Cluster_15 | Excitatory neurons_6 | 9 | 0.0014 | 3 | 0.0004 | 1005 | 0.0288 | 434 | 0.0190 | 421 | 0.0341 |
| Cluster_16 | Inhibitory neurons_3 | 48 | 0.0073 | 12 | 0.0018 | 1060 | 0.0304 | 444 | 0.0194 | 306 | 0.0248 |
| Cluster_17 | Excitatory neurons_7 | 1 | 0.0002 | 3 | 0.0004 | 780 | 0.0224 | 581 | 0.0254 | 277 | 0.0225 |
| Cluster_18 | Inhibitory neurons_4 | 36 | 0.0055 | 33 | 0.0049 | 559 | 0.0160 | 320 | 0.0140 | 137 | 0.0111 |
| Cluster_19 | Excitatory neurons_8 | 1 | 0.0002 | -- | -- | 313 | 0.0090 | 230 | 0.0100 | 122 | 0.0099 |
| Cluster_20 | Oligodendrocyte_6 | 408 | 0.0624 | -- | -- | 1 | 0.00003 | -- | -- | -- | -- |
| Cluster_21 | Excitatory neurons_9 | 1 | 0.0002 | -- | -- | 211 | 0.0060 | 133 | 0.0058 | 70 | 0.0057 |
| Cluster_22 | Endothelial | 79 | 0.0121 | 44 | 0.0066 | 148 | 0.0042 | 74 | 0.0032 | 58 | 0.0047 |
| cluster_23 | Other | 2 | 0.0003 | 5 | 0.0007 | 104 | 0.0030 | 39 | 0.0017 | 24 | 0.0019 |
| total |  | 6541 |  | 6673 |  | 34894 |  | 22900 |  | 12331 |  |

blue: less than the lower of 99% confidence interval; red: greater than the upper of 99% confidence interval

**Table 5.**
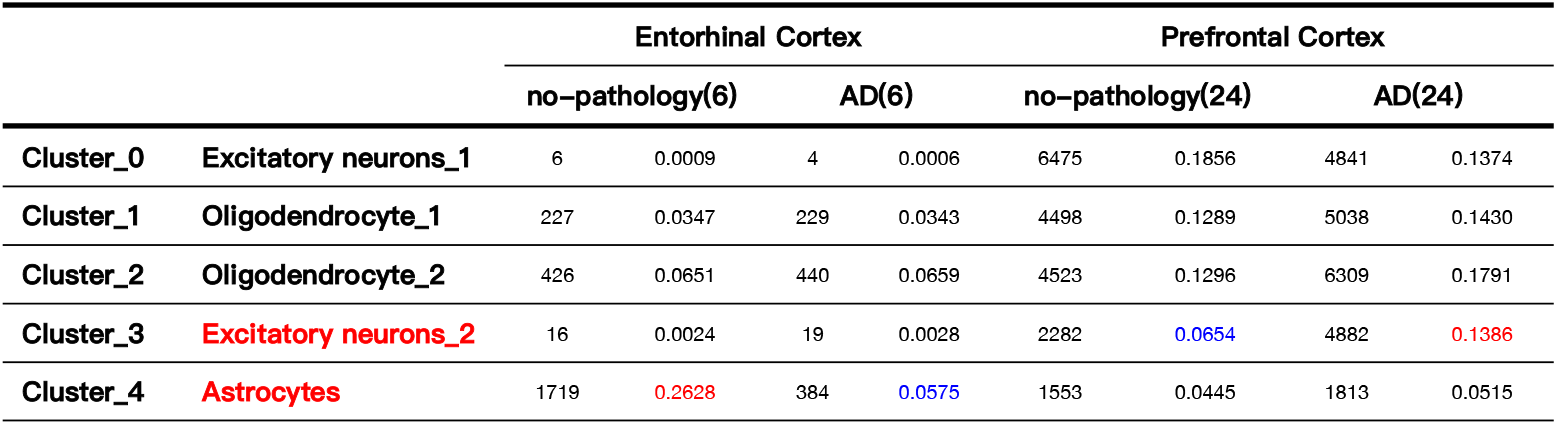
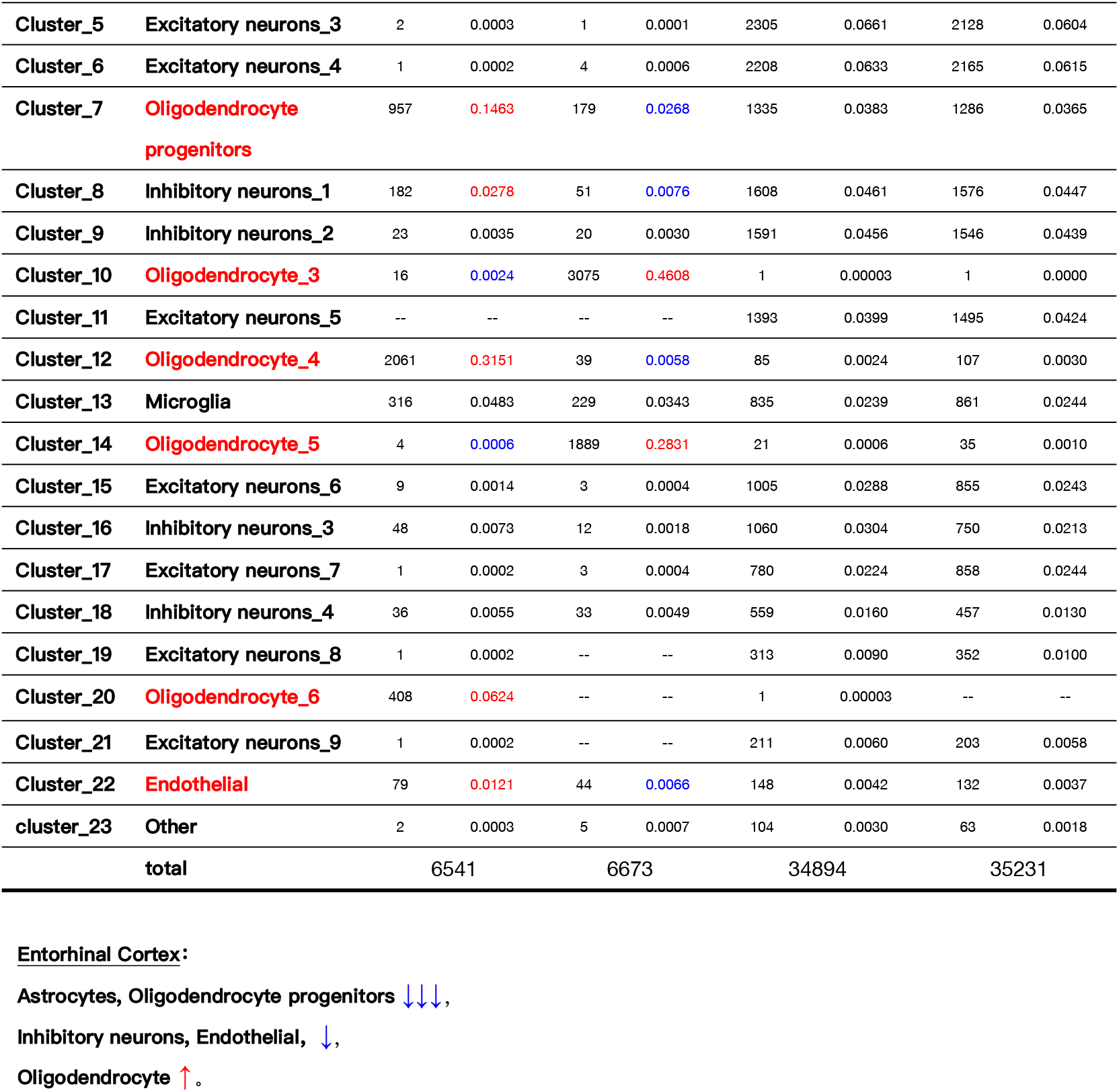
The proportion of cells for the sample in each cluster(cell sub-population.

|  |  | Entorhinal Cortex |  |  |  | Prefrontal Cortex |  |  |  |
| --- | --- | --- | --- | --- | --- | --- | --- | --- | --- |
|  |  | no-pathology(6) |  | AD(6) |  | no-pathology(24) |  | AD(24) |  |
| Cluster_0 | Excitatory neurons_1 | 6 | 0.0009 | 4 | 0.0006 | 6475 | 0.1856 | 4841 | 0.1374 |
| Cluster_1 | Oligodendrocyte_1 | 227 | 0.0347 | 229 | 0.0343 | 4498 | 0.1289 | 5038 | 0.1430 |
| Cluster_2 | Oligodendrocyte_2 | 426 | 0.0651 | 440 | 0.0659 | 4523 | 0.1296 | 6309 | 0.1791 |
| Cluster_3 | Excitatory neurons_2 | 16 | 0.0024 | 19 | 0.0028 | 2282 | 0.0654 | 4882 | 0.1386 |
| Cluster_4 | Astrocytes | 1719 | 0.2628 | 384 | 0.0575 | 1553 | 0.0445 | 1813 | 0.0515 |
| Cluster_5 | Excitatory neurons_3 | 2 | 0.0003 | 1 | 0.0001 | 2305 | 0.0661 | 2128 | 0.0604 |
| Cluster_6 | Excitatory neurons_4 | 1 | 0.0002 | 4 | 0.0006 | 2208 | 0.0633 | 2165 | 0.0615 |
| Cluster_7 | Oligodendrocyte progenitors | 957 | 0.1463 | 179 | 0.0268 | 1335 | 0.0383 | 1286 | 0.0365 |
| Cluster_8 | Inhibitory neurons_1 | 182 | 0.0278 | 51 | 0.0076 | 1608 | 0.0461 | 1576 | 0.0447 |
| Cluster_9 | Inhibitory neurons_2 | 23 | 0.0035 | 20 | 0.0030 | 1591 | 0.0456 | 1546 | 0.0439 |
| Cluster_10 | Oligodendrocyte_3 | 16 | 0.0024 | 3075 | 0.4608 | 1 | 0.00003 | 1 | 0.0000 |
| Cluster_11 | Excitatory neurons_5 | -- | -- | -- | -- | 1393 | 0.0399 | 1495 | 0.0424 |
| Cluster_12 | Oligodendrocyte_4 | 2061 | 0.3151 | 39 | 0.0058 | 85 | 0.0024 | 107 | 0.0030 |
| Cluster_13 | Microglia | 316 | 0.0483 | 229 | 0.0343 | 835 | 0.0239 | 861 | 0.0244 |
| Cluster_14 | Oligodendrocyte_5 | 4 | 0.0006 | 1889 | 0.2831 | 21 | 0.0006 | 35 | 0.0010 |
| Cluster_15 | Excitatory neurons_6 | 9 | 0.0014 | 3 | 0.0004 | 1005 | 0.0288 | 855 | 0.0243 |
| Cluster_16 | Inhibitory neurons_3 | 48 | 0.0073 | 12 | 0.0018 | 1060 | 0.0304 | 750 | 0.0213 |
| Cluster_17 | Excitatory neurons_7 | 1 | 0.0002 | 3 | 0.0004 | 780 | 0.0224 | 858 | 0.0244 |
| Cluster_18 | Inhibitory neurons_4 | 36 | 0.0055 | 33 | 0.0049 | 559 | 0.0160 | 457 | 0.0130 |
| Cluster_19 | Excitatory neurons_8 | 1 | 0.0002 | -- | -- | 313 | 0.0090 | 352 | 0.0100 |
| Cluster_20 | Oligodendrocyte_6 | 408 | 0.0624 | -- | -- | 1 | 0.00003 | -- | -- |
| Cluster_21 | Excitatory neurons_9 | 1 | 0.0002 | -- | -- | 211 | 0.0060 | 203 | 0.0058 |
| Cluster_22 | Endothelial | 79 | 0.0121 | 44 | 0.0066 | 148 | 0.0042 | 132 | 0.0037 |
| cluster_23 | Other | 2 | 0.0003 | 5 | 0.0007 | 104 | 0.0030 | 63 | 0.0018 |
| total |  | 6541 |  | 6673 |  | 34894 |  | 35231 |  |

### 3.3 Differential expression genes partition

For each pre-cluster, differentially expressed genes (DEGs) were detected using the variance-adjusted t-test (wilcoxon/logreg) as implemented in the function rank_genes_groups in SCANPY. The top 500 ranking genes were extracted for each sub-cluster, the overlap genes of t-test, wilcoxon, and logreg were identified as the DEGs for the sub-cluster.

We performed DEG enrichments for each sub-cluster using Metascape(http://metascape.org/) online tool. Then the significantly enrichment Gene Ontology (GO) and Kyoto Encyclopedia of Genes and Genomes (KEGG) pathways were determined based on the DEGs of sub-cell types, with the p-value < 0.05 as significantly enriched. The enrichment annotations for each cell type are shown in Figure 3. The shared enrichment annotations for excitatory neuron are GO:0007610: behavior, GO:0050808: synapse organization, R-HSA-112316: Neuronal System; the shared enrichment annotations for inhibitory neuron are GO:0042391: regulation of membrane potential, GO:0050804: modulation of chemical synaptic transmission, GO:0050807: regulation of synapse organization; the shared enrichment annotations for oligodendrocyte are GO:0031175: neuron projection development, GO:0034330: cell junction organization; the shared enrichment annotations for astrocyte are GO:0000902: cell morphogenesis, GO:0007420: brain development, GO:0034330: cell junction organization; the shared enrichment annotations for microglia are GO:0050778: positive regulation of immune response, WP3945: TYROBP causal network in microglia, GO:0050865: regulation of cell activation; the shared enrichment annotations for OPC are GO:0010975: regulation of neuron projection development, GO:0031175: neuron projection development, GO:0007610: behavior; finally, the shared enrichment annotations for endothelial are GO:0048514: blood vessel morphogenesis, GO:0003013: circulatory system process, GO:0061061: muscle structure development.

**Figure 3.**
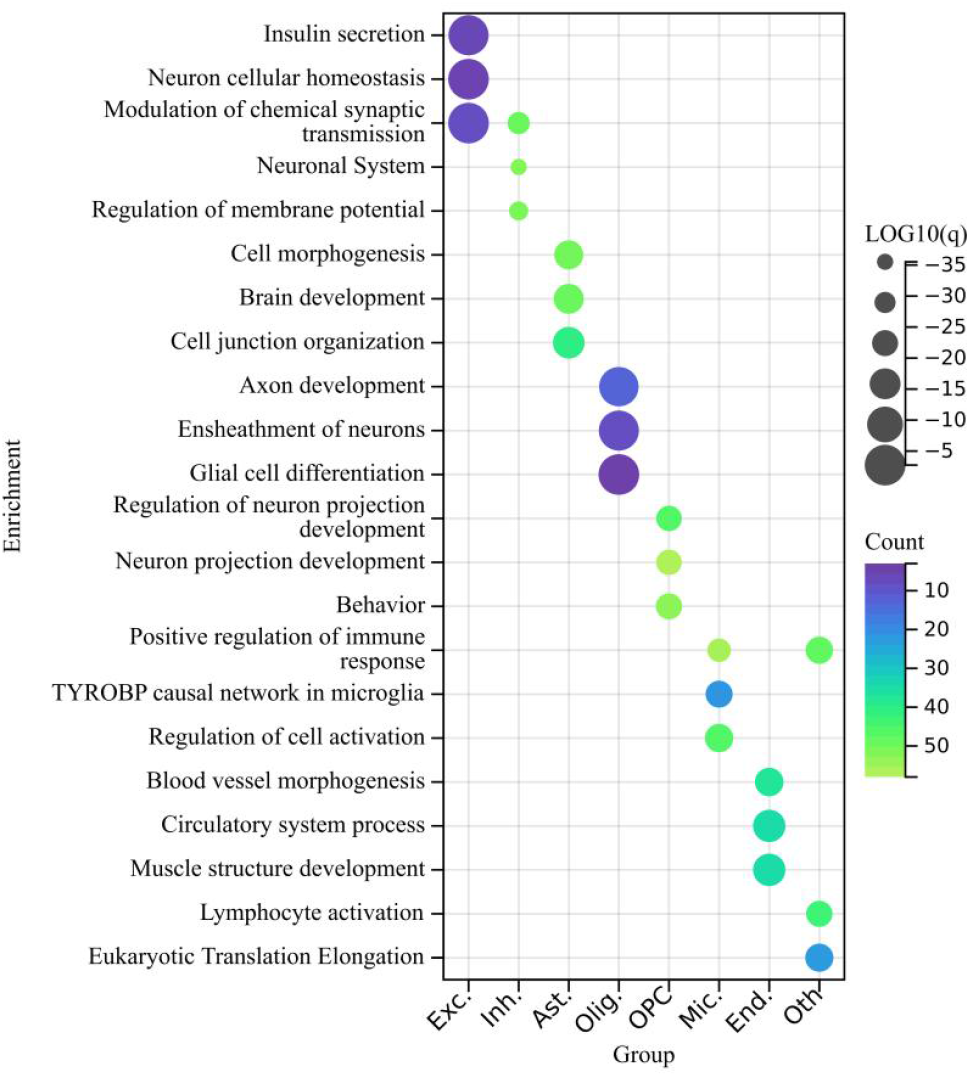
Enrichment annotations for the differentially expressed genes of each cell type.

### 3.4 Gene expression programs using cNMF

Identifying cellular process gene expression programs underlying both sub-cell type identify and sub-cell type specific or disease-dependent activities (e.g. life-cycle processes, responses to environmental cues) is crucial for understanding the biological and functional characteristics of cells. Although scRNA-seq can quantify transcripts in individual cells, each cell’s expression profile may be a mixture of both types of programs. In this study, gene programs underlying disease related sub-cell types were determined according to the cNMF analysis pipeline.

The number of components *k* used for cNMF was determined on the basis of balance between stability and reconstruction error. To reduce runtime and working memory requirements, the data were downsampled to 1500 highly variable genes. Applied to the disease-related clusters, such as Exc.2, Ast., OPC, Oli.3, Oli.4, Oli.5, Oli.6, cNMF inferred expected gene programs. We used the top 200 genes of each gene program to annotate the enrichment functions. The results were show in Figure 4. More results can be find in supplementary Figure 2–6.

**Figure 4.**
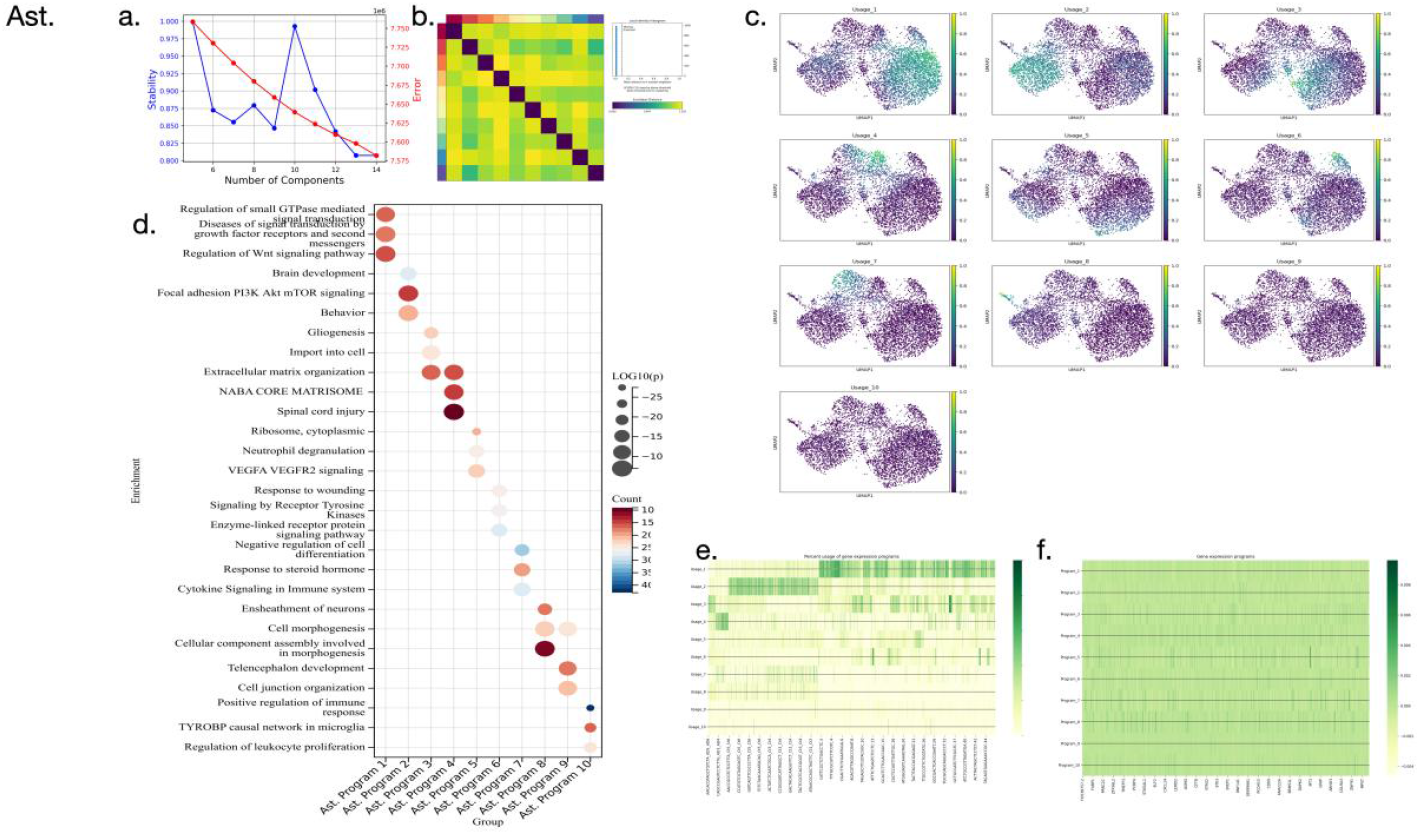
The cNMF result for sub-cluster Ast. a. the reconstruction stability and error of different component numbers. b. the consensus plot for all the program matrices. c. the normalized usages segregate on the UMAP. d. the enrichment annotations for the gene expression programs. e. the heat plot of final consensus usage matrix. f. the heat plot of final consensus program matrix.

Combine with the above findings, the function of astrocytes are cell morphogenesis, brain development, and cell junction organization. During the disease progression, regulation of small GTPase mediated signal transduction, disease of signal transduction by growth factor receptors and second messengers, regulation of Wnt signaling pathway, brain development, focal adhesion PI3K AKt mTOR signaling, behavior, gliogenesis, import into cell, extracellular into cell, and extracellular matrix organization (program 1, program 2, and program 3), were significantly affected.

## 4. Discussion

We observed that excitatory neurons, inhibitory neurons, astrocytes, and oligodendrocytes each showed distinct characteristics during the disease development, which indicates that different groups of genes respond to AD pathology in each cell type and each brain region. The AD-pathology-associated sub-clusters Exc.2, Ast. OPC, Inh.1, Oli.3, Oli.4, Oli.5, Oli.6, and End. were enriched by gene programs that have roles in protein folding and stability, neuronal and necrotic death, respectively, suggesting cell-type-specific responses to disease progression. In addition, Oli., Ast., OPCs, and End. cells in Alzheimer’s disease were enriched for genes involved in the negative regulation of cell death, indicating a coordinated response to compensate for cell stress pathways and to protect damaged cells.

Transcriptional profiling identified many differentially expressed genes in each sub-cell type of Alzheimer’s disease brain, with the most affected involving synaptic function (neurons), and signal transduction by growth factor receptors and second messengers (astrocytes).

In conclusion, this study found that, in addition to the well-studied cell-type-specific responses, that is, inflammatory responses of astrocytes and oxidative stress in neurons in Alzheimer’s disease, there is a further coordinated response that may act to boost molecular chaperone levels to protect cells against protein misfolding and that may provide additional therapeutic avenues aimed at enhancing endogenous cell type-independent responses.

## Date availability

The snRNA-seq data are available on The Rush Alzheimer’s Disease Center (RADC) Research Resource Sharing Hub at https://www.radc.rush.edu/docs/omics.htm (snRNA-seq PFC) or at Synapse https://www.synapse.org/#!Synapse:syn18485175. The ROSMAP metadata can be accessed at https://www.synapse.org/#!Synapse:syn3157322. The data are available under controlled use conditions set by human privacy regulations. To access the data, a data use agreement is needed. This registration is in place solely to ensure anonymity of the ROSMAP study participants. A data use agreement can be agreed with either Rush University Medical Center (RUMC) or with SAGE, who maintains Synapse, and can be downloaded from their websites.

## Code availability

Code used throughout this study is available upon reasonable request from the corresponding authors

## Supplementary information

Additional supporting information can be found in the supplementary figures.

---

The detailed information of participant demographics and characteristics are shown in Table 1.

**Suppl Fig. 1.**
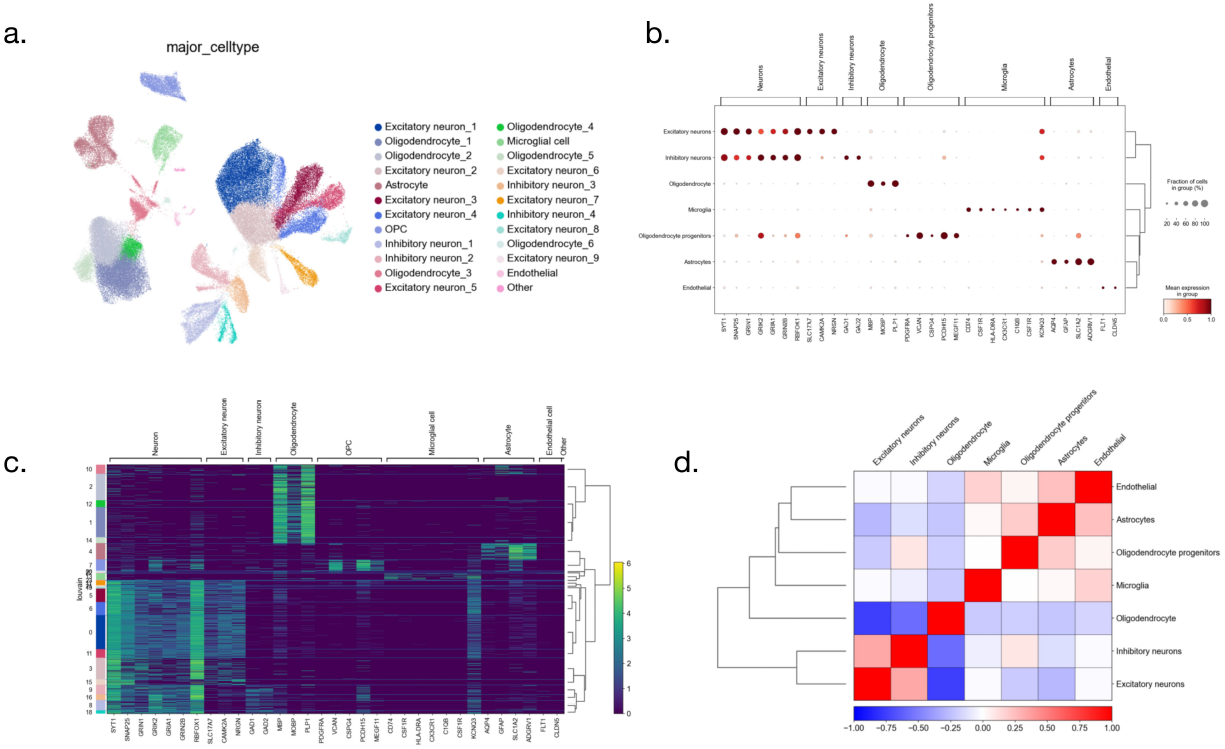
Cell type annotation and sub-clustering. a. cell type annotations for the sub-clusters. b. the expression of marker gene in each cell type. c. the hierarchical clustering heatmap of marker genes expression. d. heatmap of correlations between different cell types

**Suppl Fig. 2.**
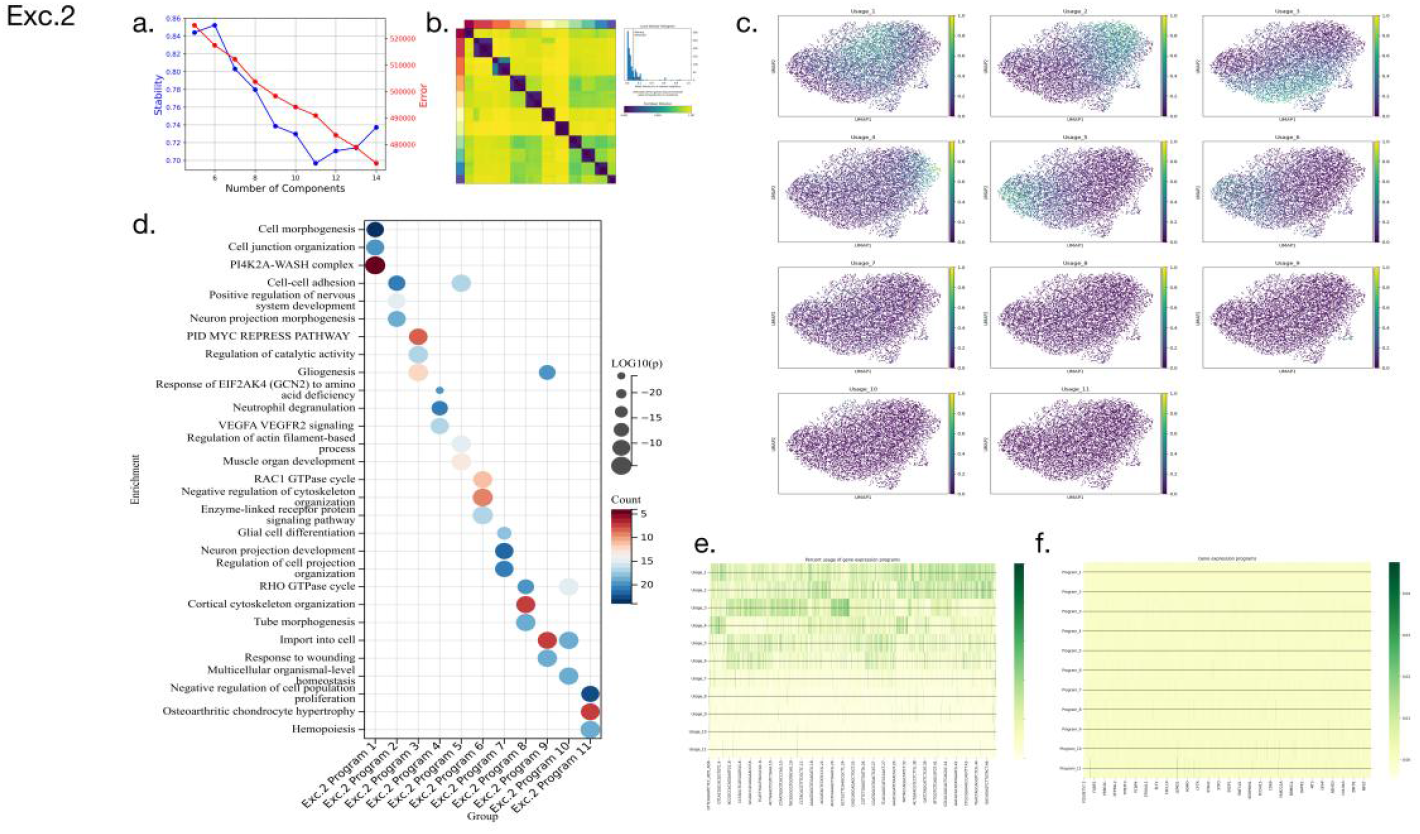
The cNMF result for sub-cluster Exc.2. a. the reconstruction stability and error of different component numbers. b. the consensus plot for all the program matrices. c. the normalized usages segregate on the UMAP. d. the enrichment annotations for the gene expression programs. e. the heat plot of final consensus usage matrix. f. the heat plot of final consensus program matrix.

**Suppl Fig. 3.**
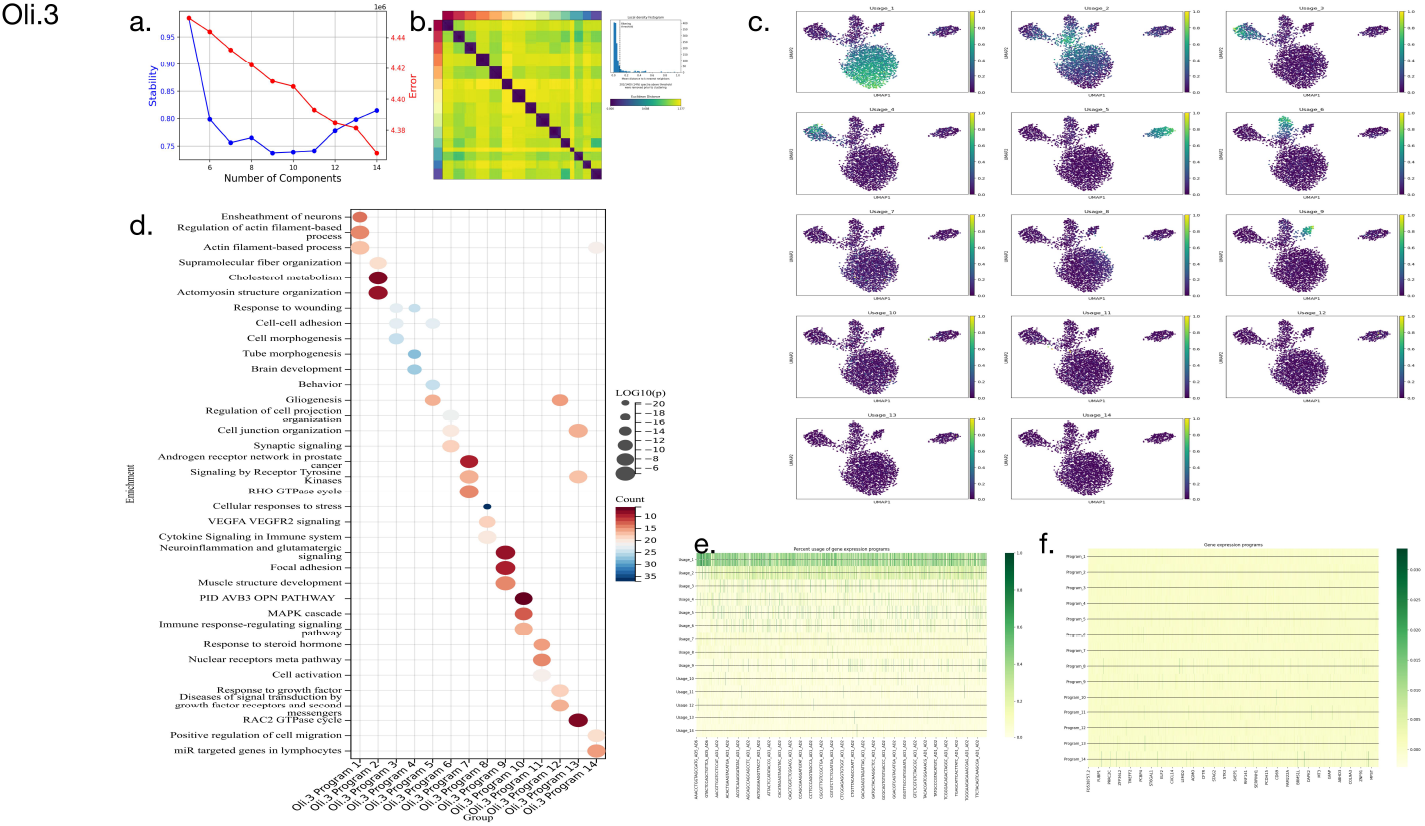
The cNMF result for sub-cluster Oli.3. a. the reconstruction stability and error of different component numbers. b. the consensus plot for all the program matrices. c. the normalized usages segregate on the UMAP. d. the enrichment annotations for the gene expression programs. e. the heat plot of final consensus usage matrix. f. the heat plot of final consensus program matrix.

**Suppl Fig. 3.**
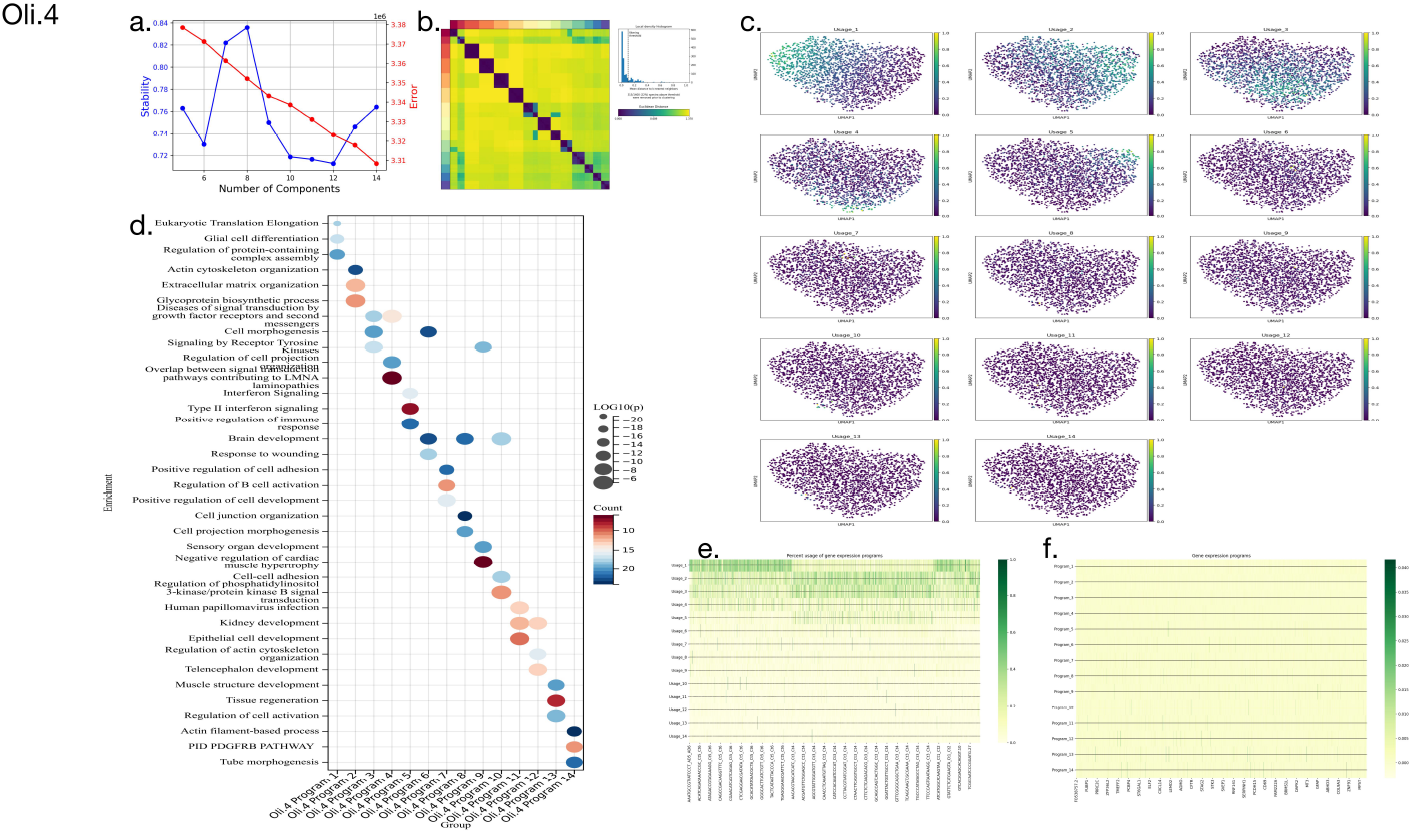
The cNMF result for sub-cluster Oli.4. a. the reconstruction stability and error of different component numbers. b. the consensus plot for all the program matrices. c. the normalized usages segregate on the UMAP. d. the enrichment annotations for the gene expression programs. e. the heat plot of final consensus usage matrix. f. the heat plot of final consensus program matrix.

**Suppl Fig. 3.**
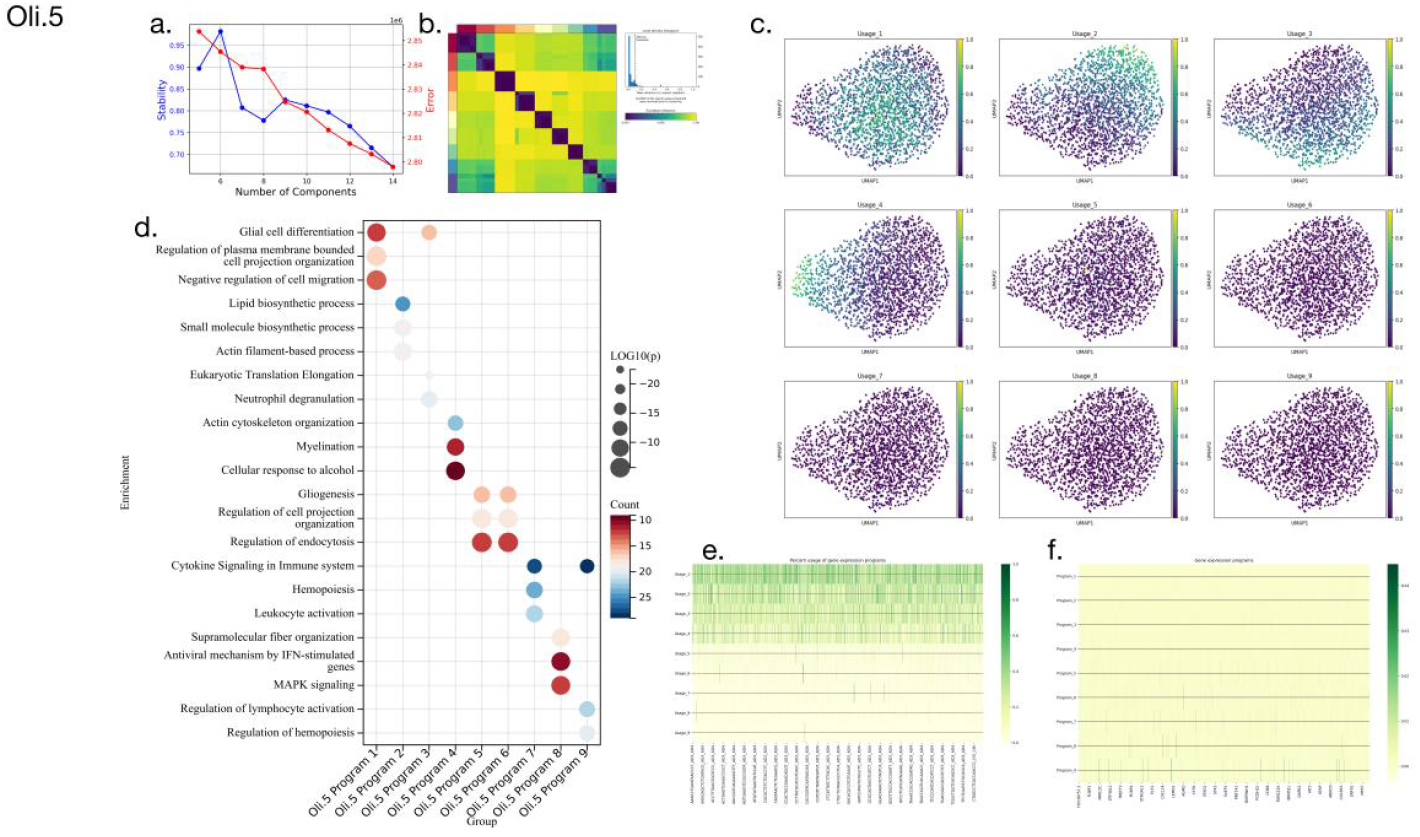
The cNMF result for sub-cluster Oli.5. a. the reconstruction stability and error of different component numbers. b. the consensus plot for all the program matrices. c. the normalized usages segregate on the UMAP. d. the enrichment annotations for the gene expression programs. e. the heat plot of final consensus usage matrix. f. the heat plot of final consensus program matrix.

**Suppl Fig. 3.**
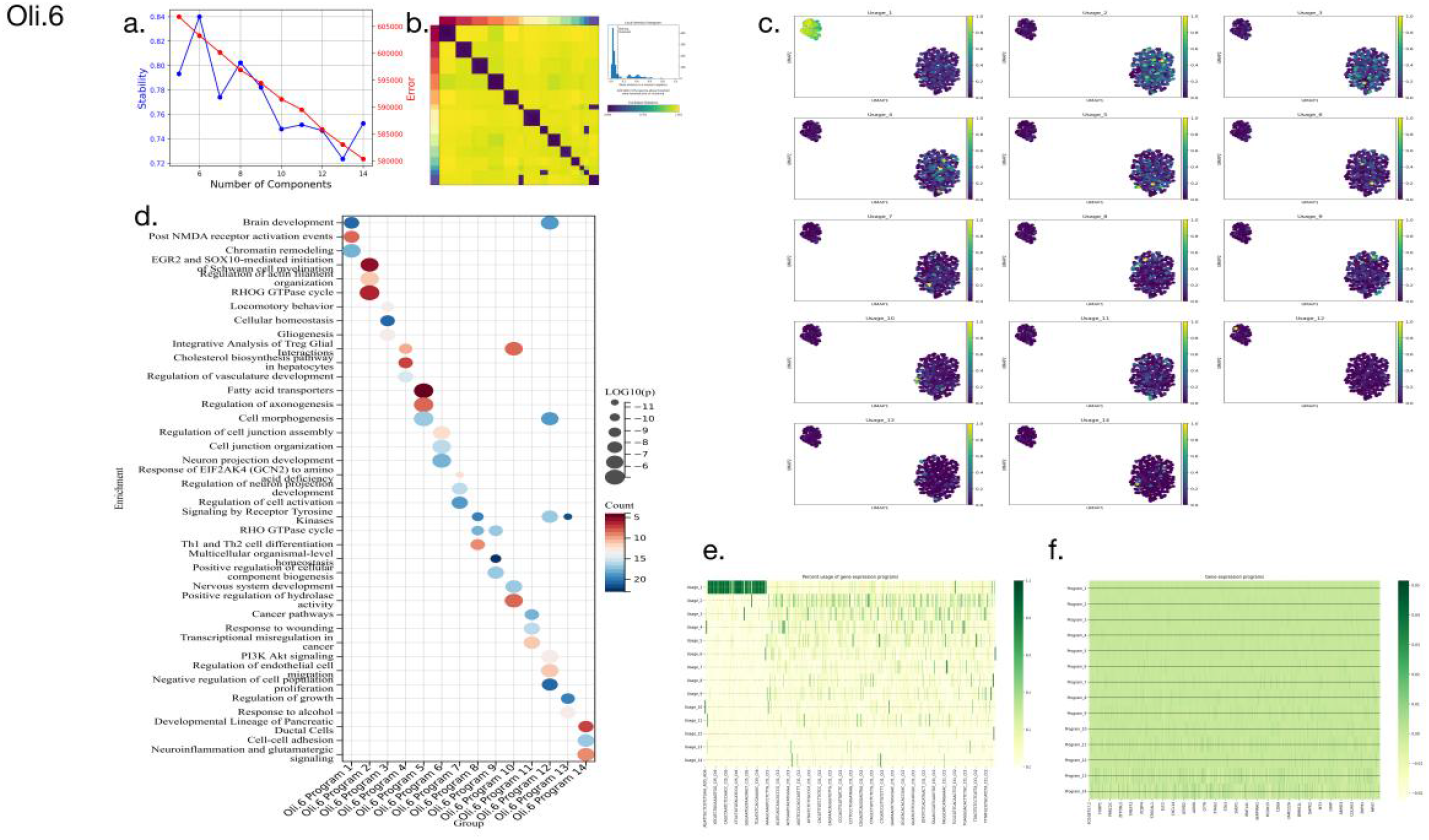
The cNMF result for sub-cluster Oli.6. a. the reconstruction stability and error of different component numbers. b. the consensus plot for all the program matrices. c. the normalized usages segregate on the UMAP. d. the enrichment annotations for the gene expression programs. e. the heat plot of final consensus usage matrix. f. the heat plot of final consensus program matrix..

**Suppl Fig. 3.**
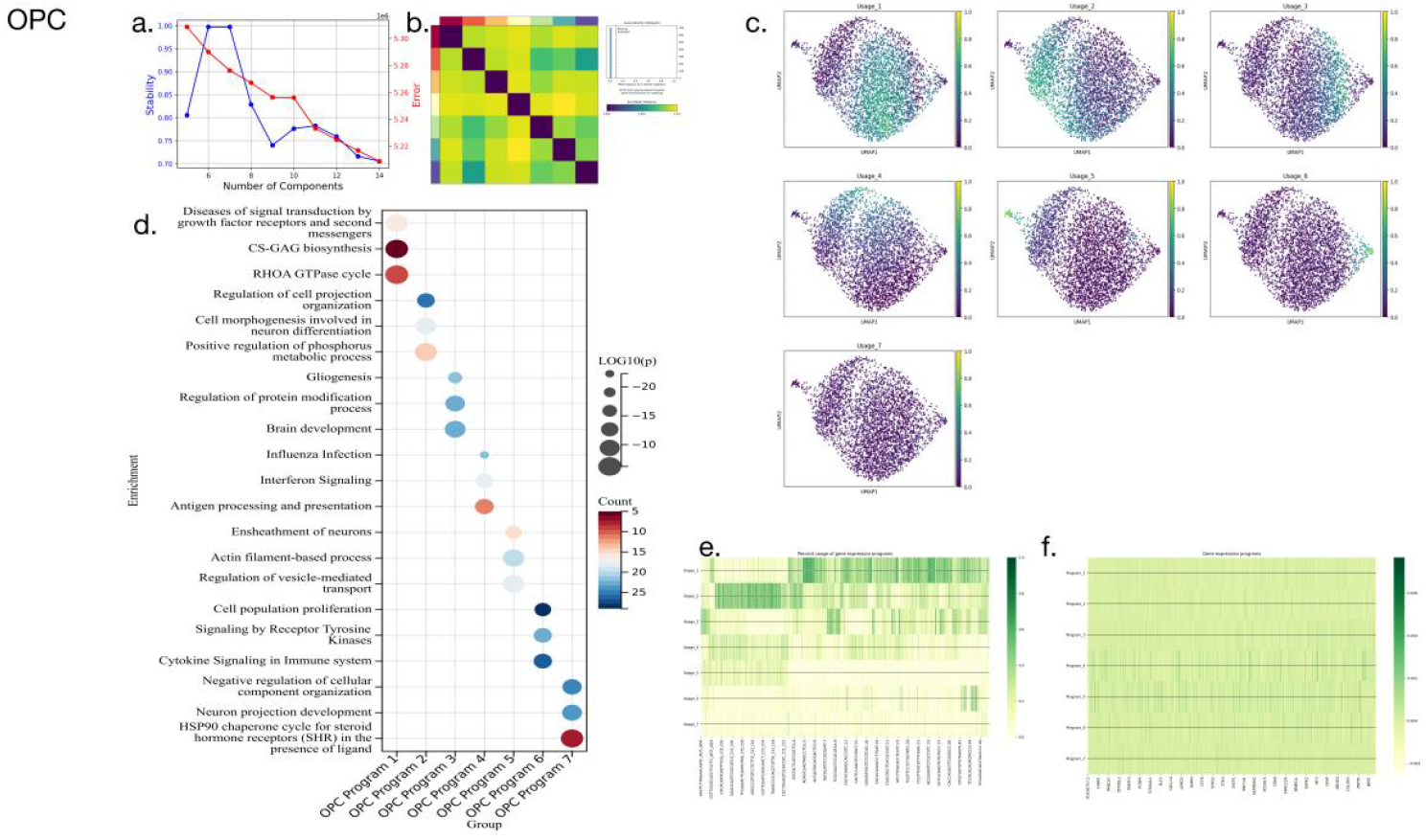
The cNMF result for sub-cluster OPC. a. the reconstruction stability and error of different component numbers. b. the consensus plot for all the program matrices. c. the normalized usages segregate on the UMAP. d. the enrichment annotations for the gene expression programs. e. the heat plot of final consensus usage matrix. f. the heat plot of final consensus program matrix.

